# Ecdysone signaling controls early embryogenesis in the short-germ hemimetabolous insect *Blattella germanica*

**DOI:** 10.1101/2022.03.10.483750

**Authors:** Josefa Cruz, Oscar Maestro, Xavier Franch-Marro, David Martín

## Abstract

Steroid hormone signaling is a central regulator of insect development especially during the post-embryonic period. However, its role during embryonic development is less understood, particularly in short-germ band hemimetabolous insects. Here, we used the short-germ band hemimetabolous insect *Blattella germanica* to analyse the functions of 20-hydroxyecdysone (20E), the most active insect steroid, signaling during the embryonic stage. We show that the heterodimeric 20E receptor, BgEcR-A and BgRXR, is present from the beginning of embryogenesis and that a stereotypical 20E-dependent cascade of nuclear receptors is first detected at the early blastoderm stage. Using parental RNAi, we show that both receptors are required for the proper formation of the germ band by controlling cell proliferation during the blastoderm stage. In addition, they are also required for the coordinated induction of the 20E-induced cascade of nuclear receptors. Finally, we show that two of these nuclear receptors, *BgHR3* and *BgFTZ-F1*, have relevant roles in germ band formation. Whereas BgHR3 controls the formation and development of the cephalic region of the germ band, *BgFTZ-F1* stripped expression suggested a role in the segmentation of the germ band. In summary, our results show that 20E-signaling is required much earlier in short-germ band hemimetabolous than in long-germ band holometabolous embryogenesis, which raises the possibility that the loss of 20E-signaling during the initial stages of embryogenesis could be a key feature in the evolution of the different types of insect development.

## Introduction

Steroid hormones play a central role in reproduction, metabolism, and development in higher organisms. In insects, they also regulate major developmental transitions, an effect that is mediated by periodic pulses of the steroid hormone 20-hydroxyecdysone (hereafter referred to as ecdysone). During post-embryonic development, these pulses promote successive molting, hence accommodating the progressive growth of the immature insect. Then, at the end of the juvenile development, a specific ecdysone burst promotes the metamorphic transition to adulthood either directly (hemimetabolous insects) or through the intermediate pupal stage (holometabolous insects) (Rewitz et al., 2013; Truman, 2019).

From a molecular perspective, ecdysone exerts its regulatory function upon binding to the Ecdysone receptor (EcR), a member of the nuclear receptor superfamily, which heterodimerizes with another nuclear receptor, the retinoid X receptor (RXR)-ortholog ultraspiracle (USP) (Henrich, 2012; Riddiford et al., 2000; Thomas et al., 1993; Yao et al., 1992; Yao et al., 1993). This heterodimer elicits then cascades of gene expression, including transcription factors *Broad-complex, E74, E93*, and a number of nuclear hormone receptors, such as *E75, HR3* and *FTZ-F1*. This core set of transcription factors mediates and amplifies the hormonal signal during each developmental transition of the insect (Cheatle Jarvela and Pick, 2017; Howard, 1995; King-Jones and Thummel, 2005; Martín, 2010; Ou and King-Jones, 2013; Riddiford et al., 2000).

In contrast to the well-characterized role of ecdysone signaling during post-embryonic transitions in hemi- and holometabolous insects, its function during embryonic development has been studied mainly in holometabolous insects, particularly in the fruitfly *Drosophila melanogaster*. Embryogenesis in *D. melanogaster* follows a long-germ band mode of development, in which almost the totality of cells of the blastoderm forms the embryo proper and all segments are formed simultaneously (Davis and Patel, 2002). From an endocrine perspective, *D. melanogaster* embryogenesis presents a single high titer pulse during mid-embryogenesis (25-50% of embryonic development), which correlates with three critical morphogenetic events: germband retraction, dorsal closure and head involution (Kraminsky et al., 1980; Maróy et al., 1988). Consistent with the timing of this ecdysone pulse, disruption of embryonic ecdysone signaling by expressing a dominant-negative EcR protein or by reducing the titer of ecdysone produces a block in the retraction of the germ band and the involution of the head (Kozlova and Thummel, 2003). Similarly, embryos derived from germline clones with *usp* mutant allele show the same phenotype (Chavoshi et al., 2010), which is indicative of EcR and USP acting together at mid-embryogenesis to promote two major morphogenetic events that shape the larval body plan. In addition to this function, ecdysone signaling is also required during mid-late embryogenesis for proper deposition of the cuticle in ectodermal tissues, such as the tracheal system and the epidermis (Ruaud et al., 2010). Ecdysone signaling, therefore, acts as a central regulator of developmental transitions in both, embryonic and post-embryonic periods in long-germ band holometabolous insects.

Contrary to the detailed information from *D. melanogaster*, very little is known about ecdysone signaling in the embryogenesis of hemimetabolous species. To gain insight into this question, here we analyzed the presence and role of ecdysone signaling in the embryogenesis of the hemimetabolous German cockroach *Blattella germanica*. Embryogenesis of *B. germanica* has been previously described, including the classification of different stages based on morphological characteristics of the embryo (Supplemental Figure S1A) (Konopová and Zrzavy’, 2005; Piulachs et al., 2010; Tanaka, 1976), and differs from that of *D. melanogaster* in some critical features. First, *B. germanica* follows a short-germ band type of embryonic development, in which only the most anterior cephalic and thoracic segments of the embryo are specified during the early syncytial blastoderm stage. The abdominal segments are then sequentially added during the germband elongation from a posteriorly located undifferentiated growth zone (GZ) of proliferating cells (Davis and Patel, 2002). Second, two clear ecdysone pulses are present during *B. germanica* embryogenesis: the first at day 6 after ootheca formation (AOF) (30–40% of embryonic development (ED)), which is associated to dorsal closure, and the second one between days 12 and 14 AOF (65–90% ED), correlating with deposition of the first instar nymphal cuticle (Supplemental Figure S1B) (Maestro et al., 2005).

Despite the presence of these two ecdysone bursts, we unexpectedly observed in previous studies that the earliest expression of ecdysone-dependent genes in *B. germanica* embryogenesis occurred around 36-48 h AOF (8-12% of embryonic development (ED) (Maestro et al., 2005; Mané-Padrós et al., 2008), which is significantly before the mid- and late ecdysone peaks. This observation suggests that, in contrast to long-germ band *D. melanogaster*, ecdysone signaling might be necessary during the very first stages of *B. germanica* embryogenesis. In the present study, we have confirmed this suggestion by showing that the heterodimeric ecdysone receptor of *B. germanica*, formed by BgEcR-A and BgRXR (Cruz et al., 2006; Martín et al., 2006), is present from the beginning of embryogenesis and that is required during the blastoderm stage for an early activation of the stereotypical ecdysone-dependent cascade of nuclear receptors and the proper formation of the germ band. Moreover, we have demonstrated that one of these nuclear receptors, BgHR3, specifically controls the formation and development of the cephalic region of the germ band, and that another nuclear receptor, BgFTZ-F1, might be involved in the regulation of germ band segmentation. Altogether, these results show that ecdysone signaling is critical during the very early stages of the short-germ hemimetabolous *B. germanica*, and reveal germ band formation as the first ecdysone-dependent transition in the life cycle of this type of insect.

## Results

### Early embryonic development in *B. germanica*

To test the possible role of ecdysone signaling during the first stages of *B. germanica* embryogenesis, we first established a detailed morphological description of the cellular dynamics during the first 60 h of embryogenesis (Figure 1A). After fertilization and fusion of the male and female pronuclei near the center of the egg, energid proliferation and migration to the periphery of the egg begins. As a result, an array of nuclei first appeared on the ventral surface of the egg at about 12 h AOF (3% ED), and then to the most dorsal part by 18 h AOF (4.4% ED). By 24 h AOF (6% ED), a uniform distribution of energids throughout the surface of the egg was achieved. Later, between 36-54 h AOF (8-13% ED), the energids located at the ventral part (0-50% of the egg width) intensively proliferate and started to migrate towards the most ventral part of the egg, where they aggregated to form a bipartite germ anlage. Finally, at 60 h AOF (15% ED), the bipartite germ anlage started to merge at the most ventral part of the egg to form the germ band (Figure 1A). At this point, the protocephalon (anterior head region) and the thorax were detectable in the germ band, as several stripes from the head segments to the T2 segment were revealed by immunostaining with the pan- segmental marker Engrailed (En) (Figure 1B). During early and late germ band elongation, 62-78 h AOF (15.1-19% ED), additional stripes of En appeared sequentially from the posterior GZ to form the abdominal part of the embryo (Figure 1B). On the other hand, cells located outside the embryonic primordium become polyploid giving rise to an extraembryonic membrane, the serosa (Supplemental Figure S2).

**Figure 1.**
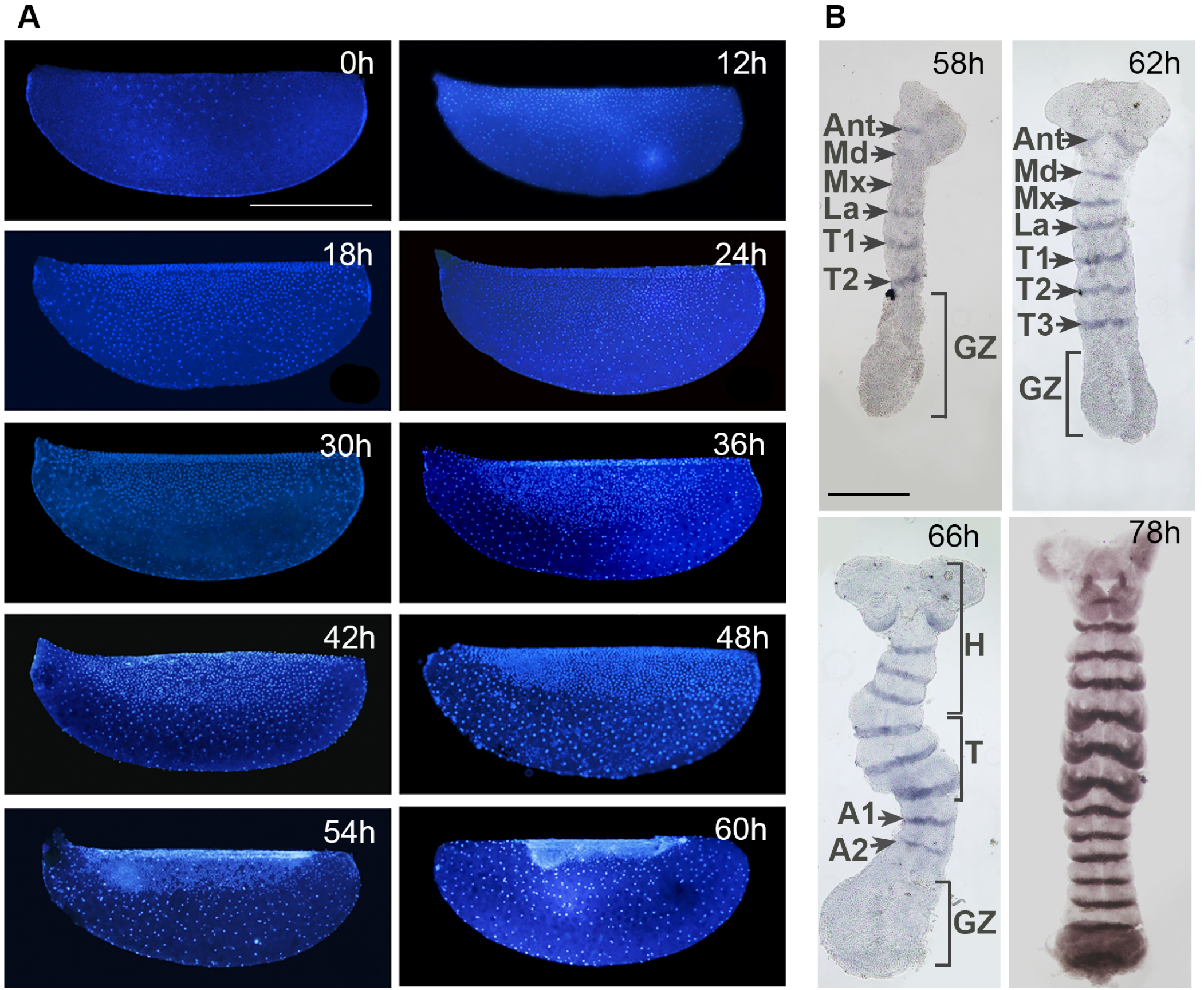
Cellular dynamics during early *B. germanica* embryogenesis. **(A)** Lateral views of DAPI-stained whole eggs during the first 60 h of embryogenesis. Time after ootheca formation (AOF) is shown by white numbers. Anterior toward the left and ventral toward the top of the eggs. After fertilization energids proliferate and migrate to the periphery of the egg (0 h AOF), first arriving at the ventral surface around 12 h AOF, and to the dorsal surface at 18 h AOF. A uniform distribution of energids is observed by 24 h AOF. Energids at the ventral part proliferate and migrate towards the most ventral part of the egg, aggregating to form a bipartite germ anlage (36-54 h AOF). The bipartite germ anlage merges at the most ventral part of the egg to form the germ band by 60 h AOF. (B) Expression of the segmental marker Engrailed during germ band formation (58-78 h AOF), showing progressive occurrence of segments in the thoracic and abdominal regions. Head (H); antennal (Ant), mandibular (Md), maxilar (Mx) and labial (La)), thoracic (T; T1-T3) and abdominal (A1-A7) segments are indicated; GZ: proliferative growth zone. Scale: 1 mm in A; 250 μm in B.

### Early occurrence of ecdysone signaling in *B. germanica* embryogenesis

To establish a correlation between the morphological changes described above and the presence of the ecdysone signaling, we examined the expression levels of the heterodimeric ecdysone receptor of *B. germanica*, BgEcR-A and BgRXR, during the period studied. As shown in Figure 2A, *BgEcR-A* and *BgRXR* receptors were maternally supplied and were present throughout the early stages of embryogenesis. Of the two *BgRXR* mRNAs described in *B. germanica* (Maestro et al., 2005), only *BgRXR-L* expression was constantly present as *BgRXR-S* was not detected until 36 h AOF. Remarkably, *in situ* hybridization revealed that at late blastoderm stage the mRNAs from both transcripts were detected surrounding the nuclei of cells that were actively proliferating and migrating to form the germ anlage (Figure 2B and C).

**Figure 2.**
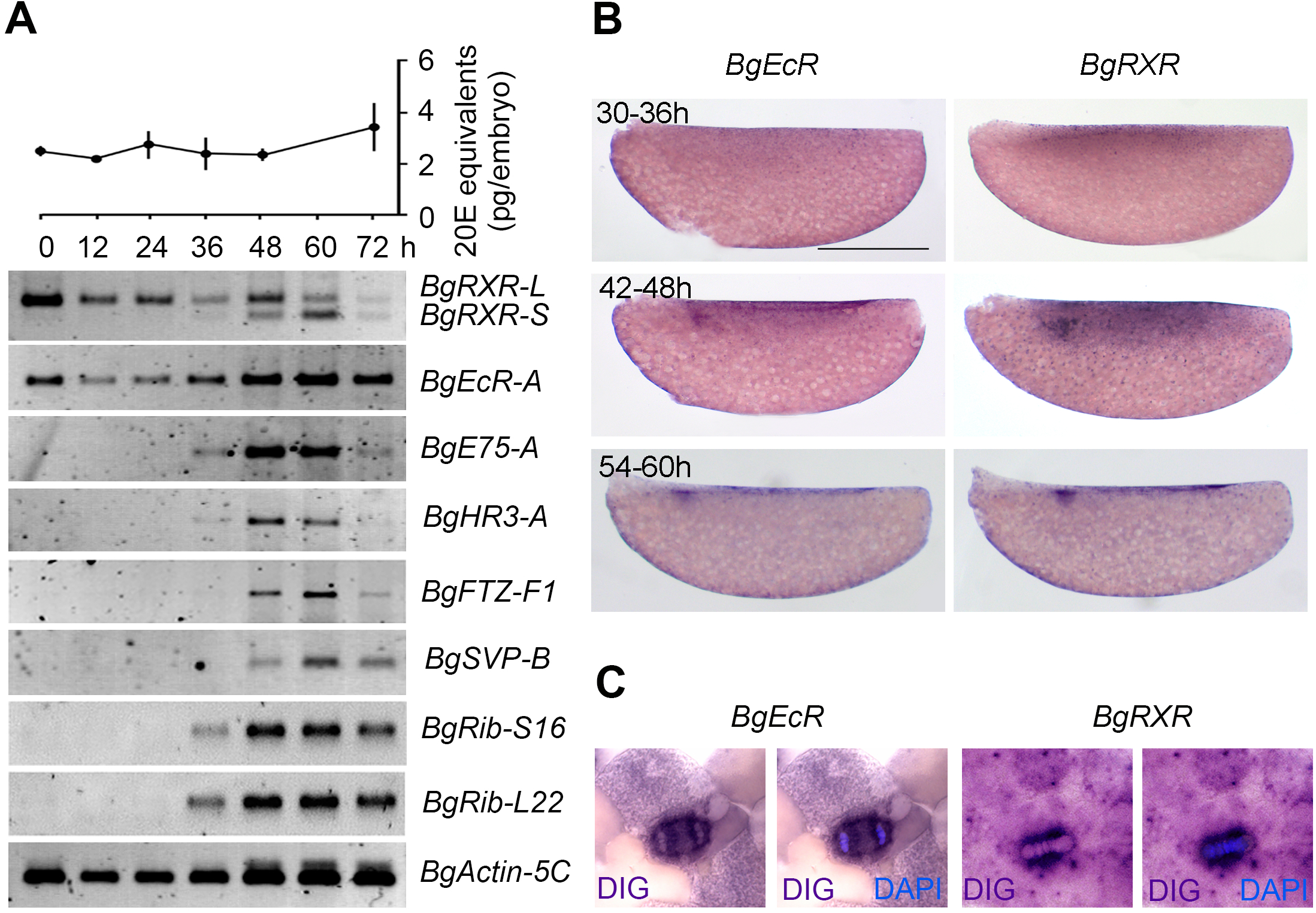
The ecdysone-dependent transcriptional cascade occurs during early embryogenesis of *B. germanica*. (A) Temporal profiles of transcription factor gene expression during early embryo development (0-60 h AOF). Equal amounts of staged embryos were analyzed by RT-PCR. *BgActin5C* levels were used as a reference. The blots are representative of five replicates. Ecdysteroid levels during the same period were measured by ELISA (upper part). (B) Expression pattern of *BgEcR-A* and *BgRXR* between 30 to 60 h AOF revealed by *in situ* hybridization. Expression of both receptors is detected in cells that proliferate and migrate to form the germ anlage. (C) *BgEcR-A* and *BgRXR* transcripts are localized around the nucleus of blastoderm cells. *in situ* hybridizations are merged with nuclear staining by DAPI. Scale: 1 mm.

Next, we analyzed the expression of the ecdysone-dependent nuclear hormone receptor genes *BgE75-A, BgHR3-A*, and *BgFTZ-F1*, which form a highly stereotypical hierarchy that is activated by ecdysone during each post-embryonic transition of *B. germanica* (Cruz et al., 2007; Cruz et al., 2008; Mané-Padrós et al., 2008; Mané-Padrós et al., 2010; Mané-Padrós et al., 2012). As Figure 2A shows, *BgE75-A, BgHR3-A* and *BgFTZ-F1* showed a transient burst of expression beginning at 36-48 h AOF, which correlates with the maternal-to-zygotic transition (Ylla et al., 2018), and ending at 60 h AOF (Figure 2A). Expression of the ecdysone-independent nuclear receptor *BgSeven-up* (*BgSvp*), and ribosomal proteins *BgRib-S16* and *BgRib-L22* was also induced by 36-48 h AOF but, in contrast to the ecdysone-dependent genes, their expression was maintained by 72 h AOF (Figure 2A). Remarkably, this early transient activation of ecdysone-dependent genes was not correlated with any detectable pulse of ecdysteroids as the levels of these hormones remained constant during the first 72 h of embryogenesis (Figure 2A). Taken together, these results show that in *B. germanica* embryogenesis ecdysone signaling activation occurs as early as 36-60 h AOF (8-12% ED), coinciding with the maternal-to-zygotic transition and the onset of proliferation and migration of the cells that will form the germ band at the most ventral part of the embryo.

### BgEcR-A and BgRXR requirement during early embryogenesis

To investigate the role of early ecdysone signaling, we use parental RNAi (pRNAi) to silence the expression of *BgEcR-A* and *BgRXR* during this period. To this aim, 5-day-old adult females of *B. germanica* were injected with *dsRNAs* encompassing the LBDs of both nuclear receptors (*BgEcRi* and *BgRXRi* animals). Specimens injected with *dsMock* were used as negative controls (*Control* animals). Under these conditions, *BgEcR-A* and *BgRXR* mRNA levels showed a marked and specific reduction at the beginning of the embryogenesis (Supplemental Figure S3A). Importantly, the removal of these receptors did not affect the proper formation of oothecae nor the levels of maternally deposited ecdysteroids (Supplemental Figure S3B and Supplemental Table S1).

Aside from 2% of unfertilized eggs, the great majority of *BgEcRi* and *BgRXRi* embryos displayed similar strong phenotypes (Figure 3A and Supplemental Table S1). As revealed by DAPI staining, they showed normal embryonic development up to 30-36 h AOF. However, from that stage, knockdown embryos presented a clear decrease in the number of cells destined to form the germinal band (Figure 3A). The reduction in cell number was not due to increased cell death (data not shown), but to impairment in cell proliferation as revealed by immunocytochemistry using anti-PH3 (Figure 3B and 3C). Despite the problems in proliferation, cells migrated correctly toward the ventral side of embryo where they condensed to form a much smaller germ band that did not progress any further (Figure 3A-A’). In contrast, absence of *BgEcR-A* and *BgRXR* did not cause the loss of the serosal cell fate as revealed by the normal density, spacing and ploidy levels of the serosal cells (Fig. 3A’), consistent with the absence of *BgEcR-A* and *BgRXR* mRNAs in these cells.

**Figure 3.**
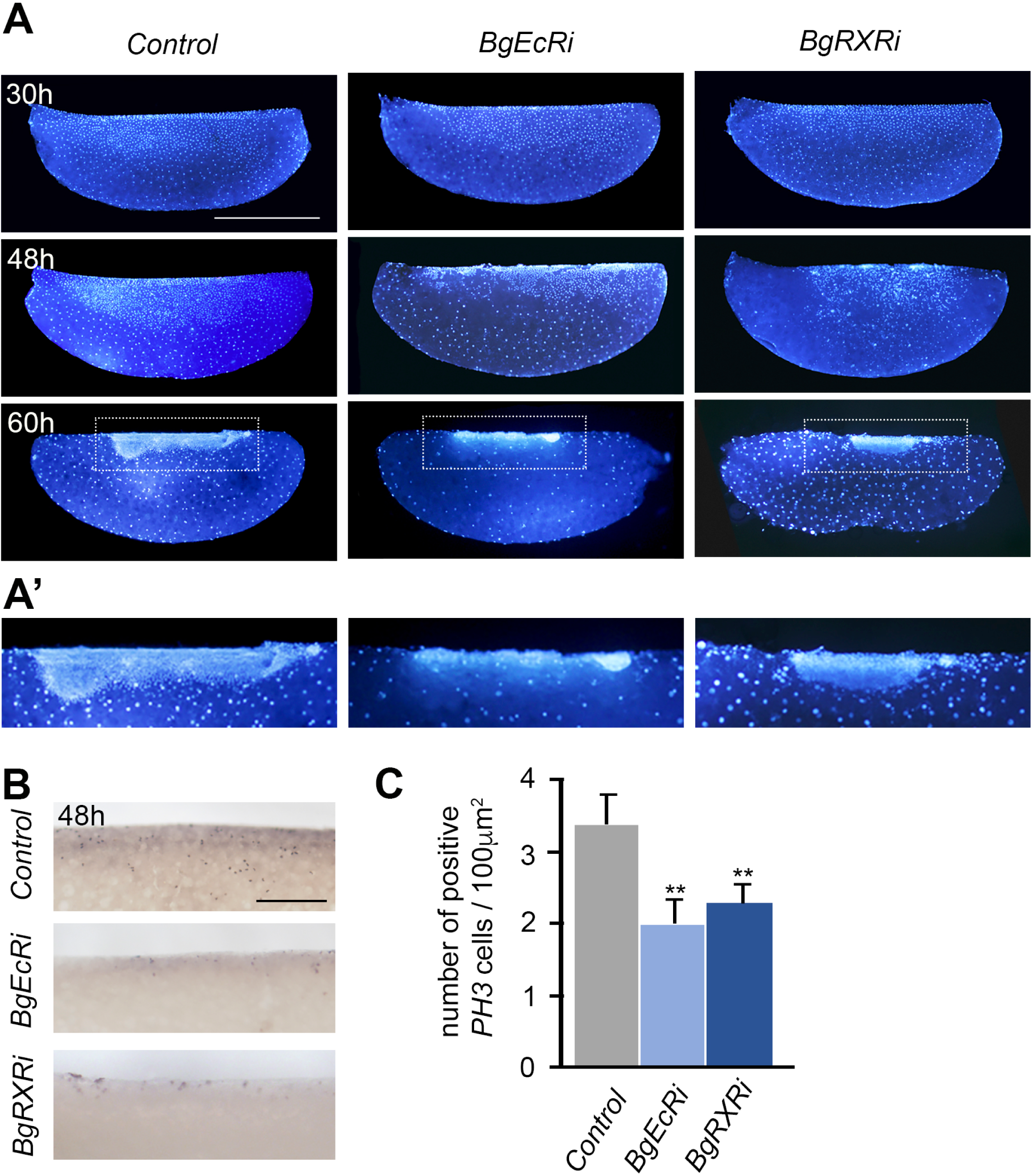
Functions of BgEcR-A and BgRXR in early embryos of *B. germanica*. (A) Development of early embryos after pRNAi of *BgEcR-A* (*BgEcRi*) and *BgRXR* (*BgRXRi*). Lateral views of DAPI-stained *Control* (left row), *BgEcRi* (middle row), and *BgRXRi* (right row) whole eggs. Time after ootheca formation (AOF) is shown by white numbers. Anterior toward the left and ventral toward the top of the eggs. Knockdown embryos presented normal embryonic development until 30-36 h AOF, but a significant decrease in the number of cells fated to become the germinal band from this point. Although the cells migrate toward the ventral part they form a significant smaller germ band that stop developing. (A’) High magnification of the rectangles in A showing the reduced germ bands of *BgEcRi* and *BgRXRi* embryos. (B) BgEcR-A and BgRXR promote cell proliferation during early embryo development. *Control, BgEcRi*, and *BgRXRi* embryos were stained with PH3 to reveal cell proliferation at 48h AOF. (C) Graph showing the average number of PH3-positive cells in *Control, BgEcRi*, and *BgRXRi* embryos. Asterisks indicate differences statistically significant as follows: p≤0.005 (**) (n=8-14). Scale bars: 1 mm in A; 200 μm in B.

Next, we analyzed if the presence of BgEcR-A and BgRXR was required for the timely activation of ecdysone-dependent genes at 36-60 h AOF. To this aim, RNA samples from staged *BgEcRi* and *BgRXRi* embryos were analyzed by qRT-PCR. As Figure 4 shows, mRNA levels of ecdysone-inducible genes *BgE75A, BgHR3* and *BgFTZ-F1* were significantly reduced in *BgEcRi* and *BgRXRi* embryos, whereas those of the ecdysone-independent gene *BgSvp-B* were not affected. Taken together, our results demonstrate that BgEcR-A and BgRXR are required during early embryogenesis in *B. germanica* for the early deployment of the ecdysone-dependent cascade of nuclear receptors as well as for the proliferation of the cells that eventually will form the germ anlage.

**Figure 4.**
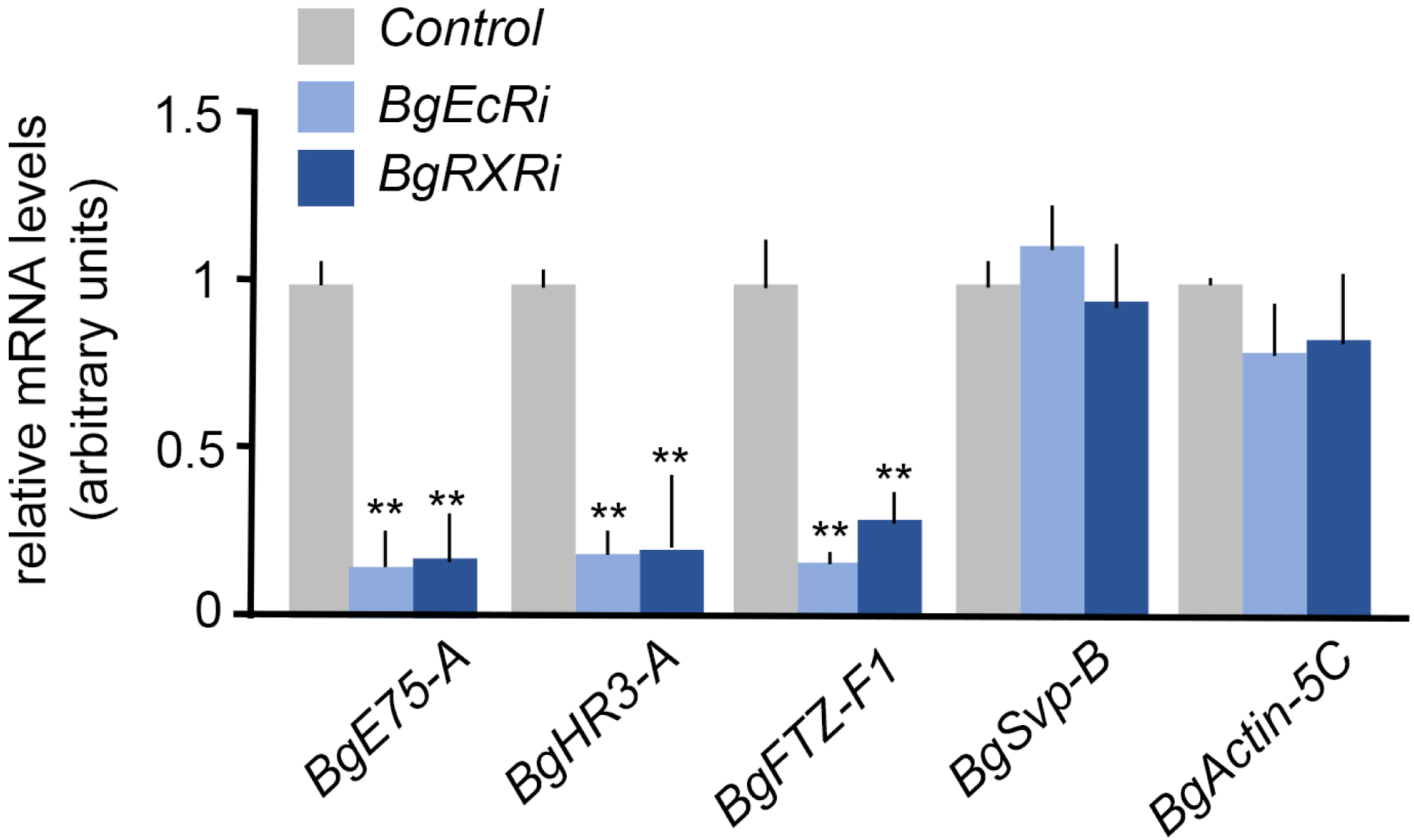
Effect of *BgEcR-A* and *BgRXR* depletion on ecdysone-dependent gene expression in early embryos of *B. germanica*. Transcript levels of nuclear receptors *BgE75-A, BgHR3-A, BgFTZ-F1* and *BgSvpB* in *Control, BgEcRi*, and *BgRXRi* embryos at 48 h AOF, measured by qRT-PCR. Fold changes in each transcript levels are relative to their expression in *Control* embryo, arbitrarily set to 1. Error bars indicate the SEM (n = 6-8). Asterisks indicate differences statistically significant compared to the control as follows: p≤0.005 (**).

### Different expression patterns of *BgHR3* and *BgFTZ-F1* during embryogenesis

The temporal coincidence between the *BgEcRi* and *BgRXRi* phenotypes and the activation of ecdysone-dependent genes suggested that components of this hierarchy might be involved in the formation and development of the germ band. To examine this possibility, we focused on the functional analysis of nuclear receptors *BgHR3-A* and *BgFTZ-F1*, as both factors have been implicated in the embryogenesis of *D. melanogaster* (Ruaud et al., 2010) and are functionally linked during postembryonic development in *B. germanica* (Cruz et al., 2008; Mané-Padrós et al., 2010). Interestingly, *in situ* hybridization analysis revealed different spatial expression patterns for both genes. Whereas *BgHR3* mRNA was strongly detected in cells that would give rise to the germ band, especially in the cephalic region (Figure 5A and A’), *BgFTZ-F1* presented a striped expression appearing from the mid-posterior half of the germ band and persisting as the germ band elongated (Figure 5B). Together, these results strongly suggest non-overlapping roles of both factors in the formation and development of the germ band during early embryogenesis of *B. germanica*.

**Figure 5.**
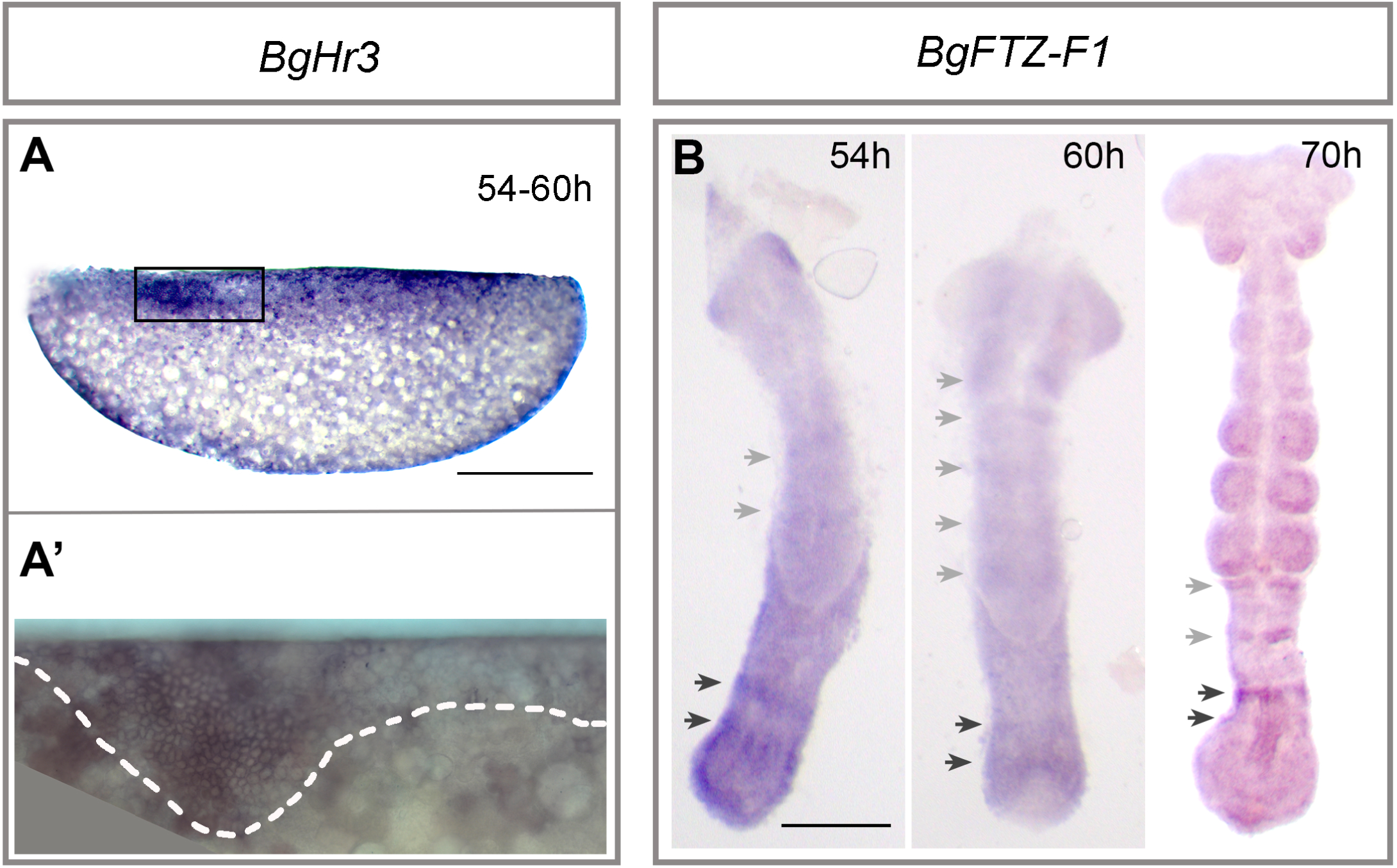
Different expression pattern of *B. germanica BgHR3* and *BgFTZ-F1* in early embryos. (A) Expression pattern of *BgHR3* during the formation of the germ band (54-60 h AOF) revealed by *in situ* hybridization. *BgHR3* transcripts are detected in cells that will form the germ band, particularly in the cephalic region (magnified in A’). (B) Striped expression pattern of *BgFTZ-F1* in the germ band (54-70 h AOF) revealed by *in situ* hybridization. *BgFTZ-F1* transcripts are detected in a striped pattern that persists as the germ band elongates. Scale bars: 1 mm in A; 250 μm in B.

### BgFTZ-F1 requirement in *B. germanica* oogenesis

The expression pattern of *BgFTZ-F1* in the GZ of the germ band pointed to a putative role in the formation of the abdomen and the segmentation process. To analyze this possibility, we carried out the same pRNAi approach that we used previously. Unfortunately, pRNAi for *BgFTZ-F1* resulted in females that did not form any ootheca regardless of the age at which the *dsBgFTZ-F1* was injected into the female, which precluded the functional analysis of the embryonic role of this factor. To ascertain the reason for this effect, we studied the expression and function of *BgFTZ-F1* in the oogenesis of adult *B. germanica* females. As Figure 6A shows, *BgFTZ-F1* was expressed in the ovary throughout the adult stage, with a modest increased towards the end of the first gonadotrophic cycle at the choriogenic stage. *BgFTZ-F1* transcripts were detected in the follicular cells that surround the basal oocyte as well as in the immature oocytes that precede the basal one, but were absent from the egg cell (Figure 6B). Administration of *dsBgFTZ-F1* in adult *B. germanica* females drastically reduced mRNA levels of *BgFTZ-F1* in 7-day-old ovaries (Supplemental Figure S3C), and resulted in a complete block of the last steps of oogenesis. A detailed analysis revealed that follicular cells from basal oocytes of *dsBgFTZ-F1*-treated females failed to rearrange the actin filaments during choriogenesis, a process that is required for proper ovulation, resulting in a completely disorganized actin structure. The cells did not fuse at the final step of oogenesis and the follicular epithelium eventually degenerated, as evidenced by the irregular distribution and aberrant shape of their nuclei (Figures 6C and D). These results demonstrate that *BgFTZ-F1* is critical for *B. germanica* oogenesis.

**Figure 6.**
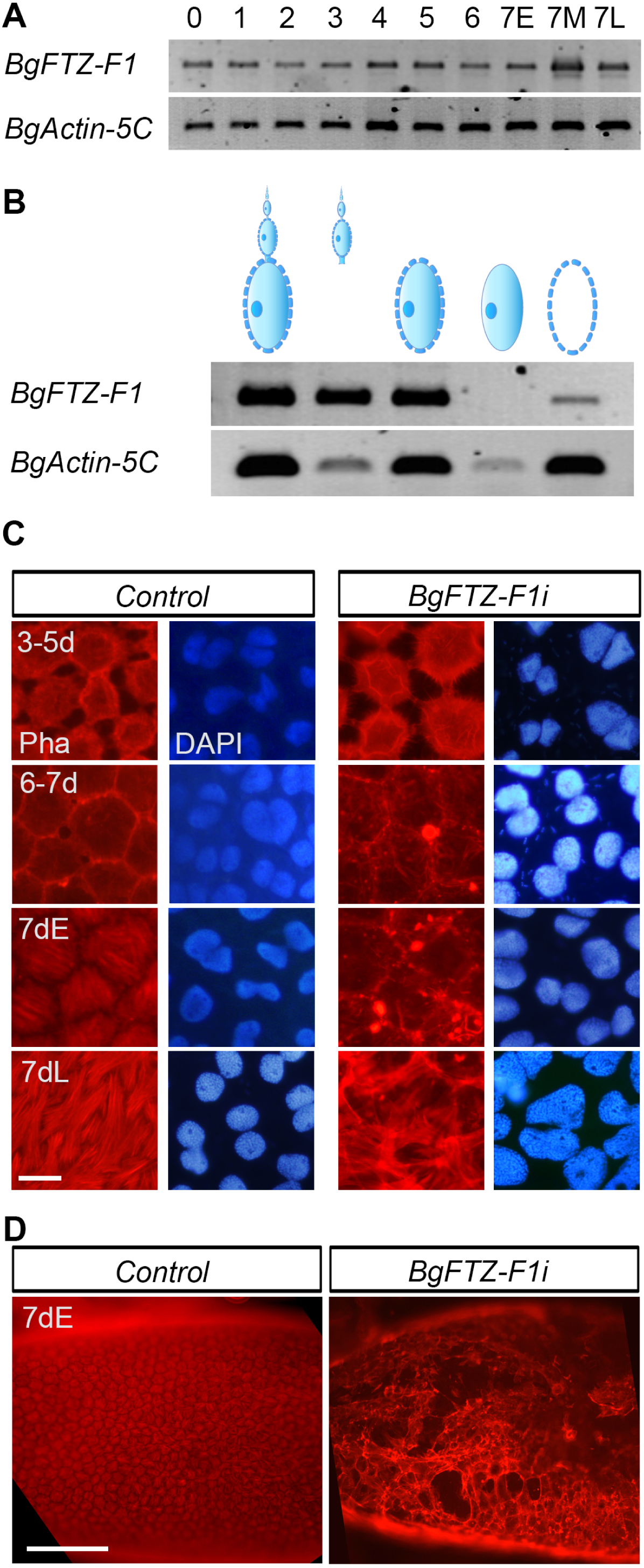
Expression and function of BgFTZ-F1 in the oogenesis of adult *B. germanica* females. (A) Spatial expression of *BgFTZ-F1* mRNA in adult ovaries analyzed by RT-PCR. Blots are representative of five replicates. Schematic representation of the different tissues analyzed (upper part). From left to right: whole ovariole; ovariole without basal oocyte; basal oocyte; basal oocyte without surrounding follicle cells; follicle cells. (B) Temporal expression (in days) of ovarian *BgFTZ-F1* mRNA during adult oogenesis. Equal amounts of total RNA from staged ovaries were analyzed by RT-PCR. 7E, 7M and 7L correspond to 7-day-old adult females with oocytes at early, mid and late stages of choriogenesis, respectively. Blots are representative of five replicates. In panels A and B, *BgActin5C* levels were used as a reference. (C) Depletion of *BgFTZ-F1* levels in adult ovaries interrupts actin filament reorganization in the follicular cells at the onset of choriogenesis. Phalloidin-tritc and DAPI stainings of follicular cells in *Control* and *BgFTZ-F1i* adult females between day 3 and oviposition. (D) Depletion of *BgFTZ- F1* in adult females results in the total degeneration of the ovarian follicle at the onset of choriogenesis revealed by Phalloidin-tritc staining. Scale bars: 50 μm in C; 500 μm in D. E: early choriogenesis; L: late choriogenesis.

### BgHR3 requirement during embryonic development

In contrast to BgFTZ-F1, pRNAi for *BgHR3* (*BgHR3i* embryos) was effective for analyzing the role of this factor during early embryogenesis. Thus, administration of *dsBgHR3* into 5-day-old adult females of *B. germanica* did not affect the formation of the ootheca and resulted in a clear decrease in embryonic *BgHR3* mRNA levels (Supplemental Figure 3D). Notably, 27% of *BgHR3i* embryos showed impaired embryonic development (Supplemental Table S2). Detailed quantification of the phenotypes indicated a graded variability within a requirement for anterior development, which was consistent with the expression of *BgHR3* in the anterior part of the germ band. We classified *BgHR3i* embryos according to the strength of the phenotype into two different classes (Supplemental Table S2). In Class I embryos only the thoracic and abdominal segments formed, although the thoracic segments developed with some truncations. The anterior cephalic region was mostly formed by undifferentiated tissue, revealed by complete or near-complete loss of En staining (Figure 7). In Class II embryos only the most anterior cephalic segments, antennae, and mandible, were missing whereas the rest of the germ band, including maxilla, labium, as well as all the thoracic and abdominal segments, developed properly (Figure 7). Altogether, our results showed that ecdysone-dependent *BgHR3* is specifically required for the proper development of the cephalic region during the early stages of *B. germanica* embryogenesis.

**Figure 7.**
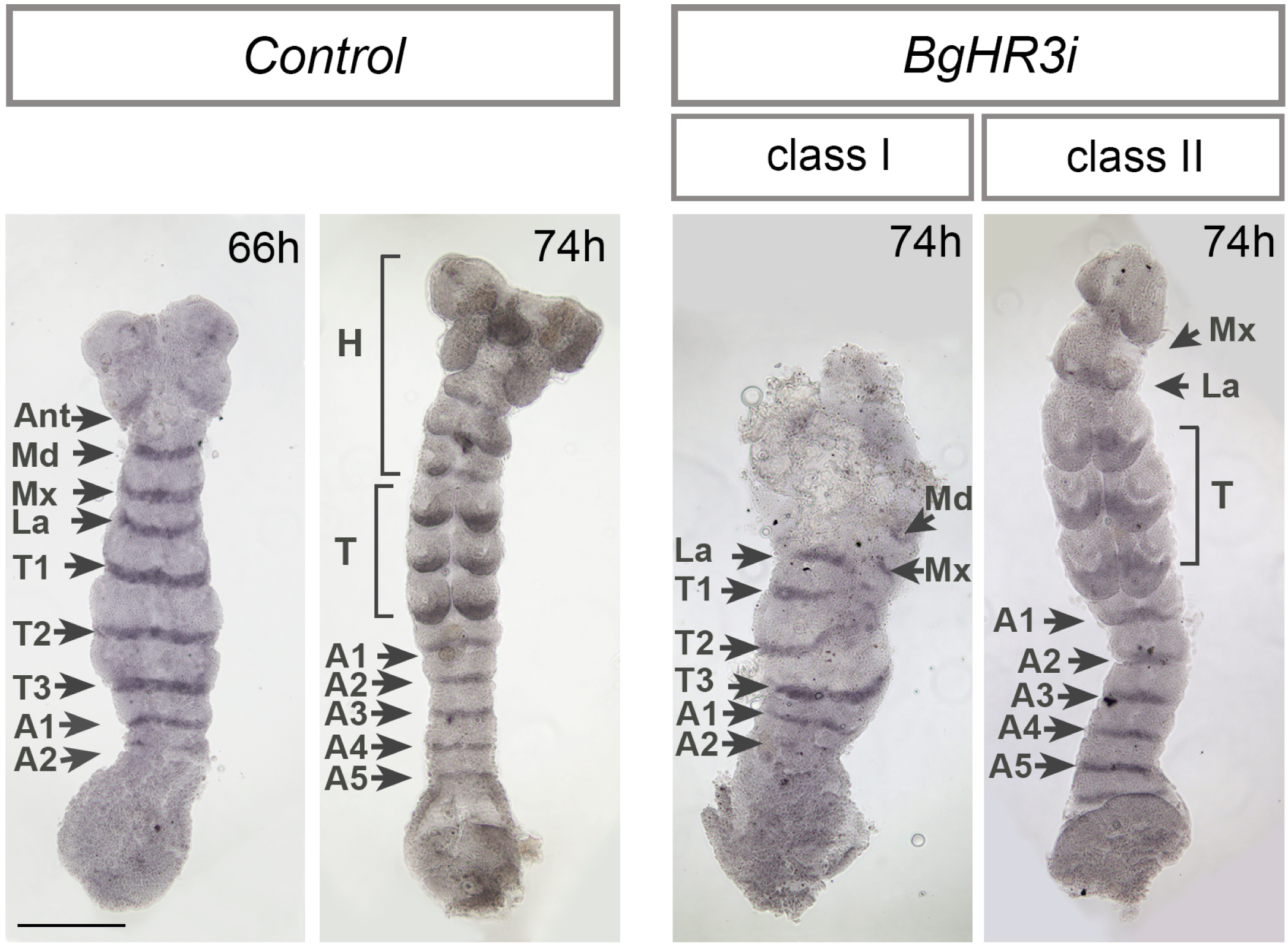
Function of BgHR3 in early embryogenesis of *B. germanica*. Development of the germ band after pRNAi of *BgHR3* (*BgHR3i*) reveals a strong impairment in the formation and development of the cephalic segments revealed by the expression of the segmental marker Engrailed. The affectation is stronger in class I *BgHR3i* germ bands compared to class II germ bands. Head (H); antennal (Ant), mandibular (Md), maxilar (Mx) and labial (La)), thoracic (T) and abdominal (A1-A5) segments are indicated. Time after ootheca formation is shown in each panel. Scale: 250 μm.

## DISCUSSION

In this study, we demonstrated that the formation and development of the germ band is the earliest ecdysone-dependent transition in hemimetabolous insect *B. germanica*. Our work shows that *BgEcR-A* and *BgRXR* are present from the beginning of embryogenesis and that they play a crucial role in controlling the first stages of the embryonic development, particularly in the proliferation of the cells that will generate the germ band. We also found that the heterodimer is also required for early deployment of the ecdysone-dependent gene cascade just at the maternal-to-zygotic transition. Finally, our findings also shown that two ecdysone dependent genes, *BgHR3* and *BgFTZ-F1* have relevant roles in germ band formation. Whereas BgHR3 controls the development of the cephalic region in the anterior region of the germ band, *BgFTZ-F1* stripped expression suggested a role in the segmentation of the germ band.

### Early ecdysone-signaling requirement in B. gemanica embryogenesis

Our results, along with previous studies, show that the ecdysone-dependent signaling cascade occurs at three different periods during *B. germanica* embryogenesis: (1) at 8-12% ED (this study); (2) at 30–40% ED, correlating with dorsal closure and deposition of the pronymphal cuticle; and (3) at 65–90% ED, which might be related with the formation of the first nymphal instar structures and the deposition of the first instar nymphal cuticle (Maestro et al., 2005; Mané-Padrós et al., 2008). The earliest period, characterized here, spanned a narrow window of time (36-60 h AOF) that starts at the maternal-to-zygotic transition and spans the formation of the germ band. Remarkably, this period is characterized by the absence of a detectable ecdysteroid pulse, although significant amounts of ecdysteroids were still detected during the first sixty hours of embryogenesis, suggesting that these hormones are maternally deposited, stored and used for the early activation of the ecdysone signaling. A similar situation is found in *Bombyx mori* embryogenesis, where ecdysone is provided not simply by *the novo* synthesis but also by dephosphorylation of maternally derived ecdysteroid conjugates (Horike and Sonobe, 1999; Makka and Sonobe, 2000; Sonobe and Yamada, 2004). Likewise, ecdysteroids required during mid-embryogenesis in *D. melanogaster* are also maternal deposited, stored as inactive conjugates in the yolk, and later converted into their active forms in an extraembryonic tissue, the amniosera (Bownes et al., 1988; Schöck and Perrimon, 2002). However, although the source of ecdysteroids at the onset of embryogenesis is the same between short- and long-germ band insects, clear differences can be observed in the ecdysteroid profiles between both types of embryogenesis. Whereas several ecdysteroid pulses are present in short-germ band insects (Bullière et al., 1979; Hoffmann et al., 1980; Lanzrein et al., 1985), a single mid-embryonic peak is observed in long-germ band insects (Maróy et al., 1988; Sullivan and Thummel, 2003; Yamada and Sonobe, 2003).

Regarding the expression patterns of *BgEcR-A* and *BgRXR-L*, our results indicate that they are maternally supplied. Interestingly, *BgRXR-L* is the only isoform detected during the first 36 h AOF, which is consistent with this isoform being the only one expressed in the ovaries of *B. germanica* during the gonadotrophic cycle (Maestro et al., 2005). Maternal contribution to embryonic *EcR* and *USP* mRNAs has been also described in *D. melanogaster* (Kozlova and Thummel, 2003; Sullivan and Thummel, 2003). Interestingly, at the blastorderm stage, *BgEcR-A* and *BgRXR* mRNAs were detected surrounding cells that were fated to become germ band, whereas were absent in cells fated to form the serosa. Consistent with their spatial expression, *BgEcR-A* and *BgRXR* loss-of-function embryos displayed impaired cell proliferation that resulted in abnormal germ bands but did not affect serosal cell fate. The control of cell proliferation by ecdysone has already been observed in the post-embryonic development of *B. germanica*, particularly in the ovarian follicle and wing cells (Herboso et al., 2015; Mané-Padrós et al., 2008). Likewise, ecdysone also controls cell proliferation in specific cell types in *D. melanogaster*. For example, the ecdysone pulse that induces the prepupal transition induces the proliferation of cells in the wing discs (Herboso et al., 2015), as well as of the abdominal histoblasts that make up the progenitor cells of the adult abdominal epidermis (Ninov et al., 2009; Verma and Cohen, 2015). However, despite the fact that ecdysone controls cell proliferation in both types of insects, our pRNAi experiments show that the requirement for ecdysone signaling is very different between the embryogenesis of short- and long-germ band insects. While BgEcR-A and BgRXR are required at the onset of embryonic development to control germ band formation, in *D. melanogaster* both receptors work together much latter, in the middle of embryogenesis, to control critical morphogenetic movements, such as the retraction of the germ band and the involution of the head, as well as for the morphogenesis of the different organs (Chavoshi et al., 2010; Kozlova and Thummel, 2003). It seems plausible, then, that the evolution from short-germ band to long-germ band was favored by the loss of control by ecdysone signaling of the initial stages of embryogenesis. These differences, therefore, might reflect a key feature in the evolution of the different types of insect embryogenesis/development. Further studies using other short- and long-germ band insects will be required to confirm this hypothesis.

### Ecdysone-dependent formation of the cephalic region through BgHR3

Our results demonstrate that the deployment of an ecdysone-dependent signaling cascade of hormone nuclear receptors at the maternal-to-zygotic transition is essential for germ band formation. The early presence of members of such cascade seems to be a conserved feature of hemimetabolan embryogenesis as it has been also observed in the hemipteran *Oncopeltus fasciatus* (Erezyilmaz et al., 2009; Surridge et al., 2011). It is important to note that this same cascade, including E75A, HR3 and FTZ-F1, is observed in all developmental transitions that are controlled by ecdysteroid pulses during the post-embryonic development of *B. germanica* (Cruz et al., 2007; Cruz et al., 2008; Mané-Padrós et al., 2008; Mané- Padrós et al., 2010; Mané-Padrós et al., 2012). Likewise, it was also found during *D. melanogaster* development, following the ecdysteroid pulses in mid-embryogenesis and between second to third larval instar and larval-pupal transitions (Sullivan and Thummel, 2003). These observations clearly indicate that this stereotypical hierarchy of ecdysone-dependent nuclear receptors acts as the temporal timer of critical developmental events throughout the life cycles of insects, and, reveal that the formation and progression of the germ band is the earliest ecdysone-dependent transition in a short-germ band type of embryogenesis.

In particular, our results have shown that nuclear receptor BgHR3 is essential for the formation and development of the cephalic region of the germ band. Interestingly, the predominant expression of *BgHR3* in the anterior region of the embryo during the blastoderm stage, before the germ band is totally formed, suggests that the spatial specification of the head by BgHR3 might precede the formation of the germ band, a possibility that has been confirmed by the reduction or elimination of the anterior head region in *BgHR3i* embryos. Interestingly, the expression and function of the cephalic gap gene *orthodenticle* (*otd*) in short-germ band insects, such as the cricket *Gryllus bimaculatus* and the beetle *Tribolium castaneum*, are consistent with the fact that head specification takes place before the onset of aggregation of the germ-anlage cells (Nakamura et al., 2010; Schröder, 2003). Therefore, our results together with previous ones, suggest that the specification of the head region in short-germ band embryos occurs at the cellularized blastoderm stage, prior to the formation of the germ band, under the control of not only cephalic gap genes (*otd*) but also of ecdysone-dependent genes (*EcR, RXR, HR3*). Whether the activation of these genes occurs in parallel or is somehow interconnected required further study.

### BgFTZ-F1 likely controls embryo segmentation and is required for adult oogenesis

Although pRNAi for *BgFTZ-F1* has failed to provide evidence for an embryonic role of this factor, clear spatial differences were observed for *BgHR3* and *BgFTZ-F1* expression. These differences strongly suggest non-overlapping functions of both receptors during early *B. germanica* embryogenesis, a situation that contrasts with that observed in post-embryonic development, where both factors overlap in regulating the same developmental processes (Cruz et al., 2007; Cruz et al., 2008). In contrast to the cephalic expression of *BgHR3*, the striped expression of *BgFTZ-F1* in the thorax and abdomen of germ bands points to a possible function in controlling segmentation. This pattern of expression seems to be conserved in short- and intermediate-germ band embryogenesis as similar *FTZ-F1* expression in pair-rule stripes has been observed also in the beetles *Dermestes maculatus* and *T. castaneum* (Heffer et al., 2013; Xiang et al., 2017), suggesting a conserved role of FTZ-F1 in segmentation. In fact, functional studies in *T. castaneum* revealed that depletion of *Tc-FTZ-F1* resulted in clear pair-rule segmentation defects (Heffer et al., 2013; Xu et al., 2010). Interestingly, the pair-rule function of FTZ-F1 is conserved in the embryogenesis of long-germ band *D. melanogaster*, although in the fruit fly this factor is expressed ubiquitously in the embryo (Yussa et al., 2001). In this case, segment specification was achieved simultaneously by the interaction of ubiquitous FTZ-F1 with the homeodomein protein Fushi Tarazu (Ftz), which was expressed in pair-rule stripes (Florence et al., 1997; Guichet et al., 1997; Yu et al., 1997). In this regard, it is interesting to note that E75A, another member of the cascade, exerts pair-rule functions in *O. fasciatus* (Erezyilmaz et al., 2009), suggesting that a combination of ecdysone-dependent nuclear receptors might be involved in the control of segmentation process in short-germ band insects. Unfortunately, no embryonic functions for E75A have been characterized in any other short-germ band insect. In our case, were not able to obtain a reliable depletion of embryonic *BgE75A* by pRNAi, probably due to the small size of the isoform-specific region (data not shown). It would be interesting in the future, then, to determine to what extend ecdysone-dependent nuclear receptors are involved in the control of segmentation to ascertain the ancestral functions of these genes in embryogenesis.

Finally, we have demonstrated a critical contribution of BgFTZ-F1 in the oogenesis of adult *B. germanica* females. *BgFtz-F1* is expressed throughout the ovary during the whole gonadotrophic cycle, and its depletion results in the complete block of oogenesis due, at least in part, to failure in the reorganization of the actin filaments of the follicle cells during choriogenesis. Interestingly, FTZ-F1 is also required for completing oogenesis in *T. castaneum* (Xu et al., 2010) and *D. melanogaster* (Beachum et al., 2021; Knapp et al., 2020). In the fly, like in *B. germanica*, this factor is detected throughout the ovary, and is required for follicle cells to differentiate properly into the final maturation stage (Beachum et al., 2021; Knapp et al., 2020). Likewise, the mammalian FTZ-F1 homolog, steroidogenic factor-1 (SF-1) has been also detected in somatic follicle cells in rodents and humans (Hinshelwood et al., 2003; Tajima et al., 2003), and its depletion in mouse granulosa cells results in the loss of developing follicles (Pelusi et al., 2008). Together, these results highlight the conservation of the critical role of FTZ-F1/SF- 1 in the control of oogenesis in insects and mammals.

In summary, our work shows the pivotal role of ecdysone signaling during early embryogenesis in the hemimetabolous *B. germanica*, and identifies the formation and development of the germ band as the first ecdysone-dependent transition in a short-germ band type of embryogenesis. We can conclude, therefore, that in short-germ band hemimetabolous insects, ecdysone signaling acts as a central regulator of developmental transitions in embryonic and post-embryonic periods. However, given that some holometabolan species also undergo short-germ band embryonic development, further research using different short-germ band holometabolan species, would be required to ascertain the early presence of ecdysone signaling and to generalize the regulatory role of this pathway in this type of embryonic development.

## MATERIALS AND METHODS

### Insects

Specimens of *B. germanica* were obtained from a colony reared in the dark at 30 ± 1 °C and 60-70 % r.h. Under our rearing conditions, embryogenesis in *B. germanica* lasts 17 days, during which eggs developed within an ootheca that was carried out by the female. All dissection and tissue sampling were carried out in Ringer’s saline using carbon dioxide-anesthetized specimens.

### Semiquantitative reverse transcriptase polymerase chain reaction (RT-PCR)

RT-PCR followed by southern blotting with specific probes was used to determine the embryonic and ovarian expression pattern of genes involved in the research. Total RNA was extracted from embryos and ovaries and cDNA synthesis was carried out as previously described (Cruz et al., 2006). Primers for the amplification of the corresponding genes can be found at Supplementary Table S3. As a reference the same cDNAs were subjected to RT-PCR with a primer pair specific for *B. germanica Actin5C*. cDNA samples were subjected to PCR with a number of cycles within the linear range of amplification for each transcript depending on the tissue and physiological stage. cDNA probes for Southern blot analyses were generated by PCR with the same primer pairs and labelled with fluorescein (Amersham Biosciences).

### Quantitative real-time reverse transcriptase polymerase chain reaction (qRT-PCR)

Total RNA extraction and cDNA synthesis was carried out as described above. Relative transcripts levels were determined by quantitative real-time PCR (qPCR), using iQ SYBR Green supermix (Bio-Rad), and forward and reverse primers (final concentration: 100nM/qPCR) in a 20 µl final volume (see Supplementary Table S3 for primer sequences). The qPCR experiments were conducted with the same quantity of embryo equivalent input for all treatments and each sample was run in duplicate using 2 µl of cDNA per reaction. All the samples were analyzed using an iCycler and iQ Real Time PCR Detection System (Bio-Rad). For each standard curve, one reference DNA sample was diluted serially. RNA expression was normalized in relation to the expression of *BgActin5C*.

### Whole-mount hybridization in situ

Digoxigenin-labeled RNA probes (sense and antisense) were generated by transcription *in vitro* using SP6 or T7 RNA polymerases (Roche) as previously described (Cruz et al., 2006). For BgEcR-A and BgHR3-A probes, 527-bp and 520- bp fragments, respectively, containing the specific A/B domains were used. For BgFTZ-F1 and BgRXR probes, 542-bp and 497-bp fragments within their ligand binding domains were used.

### Microscopy and histological analysis

For immunocytochemistry, embryos were dissected and fixed in 4% paraformaldehyde-PEM (100mM Pipes, 2mM EGTA, 1mM MgSO4), permeabilized in PBST (PBS, 0,1% Triton) and incubated overnight at 4ºC with the primary antibody diluted in PBSTBN (PBST, 0,1% BSA, 5% NGS). The following primary antibodies were used at indicated dilution: mouse anti- engrailed/invected 4D9 (Patel et al., 1989), 1:1) and rabbit anti-phospho-Histone H3 (Ser10) (Merk, 1:1000). They were then washed with PBST and incubated with the secondary antibody: anti-mouse IgG-peroxidase (Merk, 1:100) or anti-rabbit IgG-peroxidase (Merk, 1:50) diluted in PBST for to 2 h and washed with PBST. The DAB reaction was performed for 3-5 min and embryos mounted for visualization in 50% glycerol in PBS. For fluorescent imaging, dissected ovaries were dissected in Ringer and fixed in 4% paraformaldehyde in PBS. Permeabilized in PBST and incubated for 20 min in a 300ng/ml solution of phaloidin-TRITC in PBS. Rinsed with PBST and mounted in Vectashield medium with DAPI (Vector Laboratories, H1200). All samples were examined with a Zeiss Axiophot microscope, and images were subsequently processed using Adobe photoshop.

### Gene knockdown by parental RNA interference (pRNAi)

The primers used to generate templates via PCR for transcription of the dsRNAs are described in Supplementary Table S3. Synthesis of double-stranded RNA was performed as previously described (Cruz et al., 2006). A volume of 1 µl of each dsRNA solution (3-5 µg/µl) was injected into the abdomen of adult females. Under our rearing conditions, the first adult gonadotrophic cycle lasts seven days. During this period, only the basal oocyte of each ovariole matures and growths, whereas the remaining oocytes of the ovariole keep immature. When maturation is completed, basal oocytes form the chorion and are oviposited into an attached ootheca where embryogenesis takes place. To determine the optimal range of adult female age for pRNAi experiments, we injected *dsRNAs* into adult females from day 0 to day 7, and found that injection into 5-days-old females produced the best pRNAi effect for *BgEcR-A, BgRXR* and *BgHR3*. In the case of *BgFTZ-F1*, no oothecae were formed regardless of the female age at which the *dsRNA* was injected. For the analysis of the role of BgFTZ-F1 in oogenesis, *dsRNA* for *BgFTZ- F1* was injected into newly ecdysed adult females. *Control dsRNA* (*Control*) consisted of a non-coding sequence from the pSTBlue-1 vector.

### Quantification of embryonic ecdysteroid titers

20 staged oothecas for each time point were preserved in 500ul of methanol, homogenized and centrifuged (10 min at 18000 x g). The remaining tissue was re-extracted twice in 0.5 mL methanol and the resulting methanol supernatants were dried using a SpeedVac. Samples were resuspended in enzyme immunoassay buffer (0.4cM NaCl, 1cmM EDTA, 0.1% bovine serum albumin in 0.1 M phosphate buffer). ELISA was performed according to the manufacturer’s instructions using a commercial ELISA kit (Bertin Bioreagents) that detects ecdysone and 20- hydroxyecdysone with the same affinity. Absorbance was measured at 405 nm on a plate reader, SpectramaxPlus (Molecular Devices, Sunnyvale, CA) using GraphPad Prism v4.00 software (GraphPad Software Inc., San Diego, CA).

## Acknowledgements

Support for this research was provided by the Spanish MINECO (Grants CGL2014-55786-P and PGC2018-098427-B-I00 to D.M. and X.F-M.) and by the Catalan Government (2014 SGR 619 to D.M. and X.F-M.). The research has also benefited from FEDER funds.

## Author Contributions

Conception and design of the project was done by J.C. and D.M. J.C. and O.M. performed the experiments. The analysis of the data was conducted by J.C. O.M. X.F.-M. and D.M. D.M. wrote the manuscript and J.C and X.F.-M. revised the manuscript. All authors approved the final manuscript.

